# Diversifying mechanisms in the on-farm evolution of crop mixtures

**DOI:** 10.1101/009829

**Authors:** Mathieu Thomas, Stéphanie Thépot, Sophie Jouanne-Pin, Nathalie Galic, Carine Remoué, Isabelle Goldringer

## Abstract

While modern agriculture relies on genetic homogeneity in the field, some farmers grow genetically heterogeneous crops and exchange seeds. Such diversifying practices associated with seed recycling may allow adaptation of crops to their environment. This socio-genetical model constitutes an original experimental evolution design called On-Farm Dynamic Management (OFDM). Studying OFDM can help understanding how evolutionary mechanisms shape crop diversity submitted to diverse agro-environmental conditions. We studied a farmer-led initiative where a mixture of four French wheat landraces called “Mélange de Touselles” (MDT) was created and distributed within a farmers’ network. Fifteen populations derived by farmers from the initial mixture were sampled after 2 to 7 generations of cultivation on their farm. Twenty-one space-time samples of 80 individuals were genotyped using 17 microsatellites markers and characterized for their heading date in a “common-garden” experiment. Gene polymorphism was studied using four markers located in earliness genes. An original network-based approach was developed to depict the particular and complex genetic structure of the landraces composing the mixture. A rapid differentiation of the mixture was detected, larger at the phenotypic and gene levels compared to the neutral genetic level, indicating a potential divergent selection. We identified two interacting selection processes: variation of the mixture component frequencies and evolution of the within-variety diversity, that shaped the standing variability available within the mixture. These results confirm that farmers’ practices increase genetic diversity and allow crop evolution, which is critical in the context of global change. OFDM appears as a promising model of crop experimental evolution.

## 1 Intro

## 2 Introduction

Genetic diversity is assumed to be of major importance for the adaptation of both wild and cultivated species to future environmental changes (Barrett and Schluter, 2008; Mercer and Perales, 2010). As farmers around the world utilize various farming practices and grow different species for different uses in different agroecosystems, they *de facto* contribute to the on-farm dynamic management of crop diversity. On-farm dynamic conservation of agrobiodiversity, is a complementary strategy to *ex situ* conservation, that allows genetic resources to continuously adapt to changing environments (Bretting *et al.*, 1997; Maxted *et al.*, 1997; Negri and Tiranti, 2010; Enjalbert *et al.*, 2011). While in Dynamic Management (DM), crop populations are mainly submitted to natural selection in experimental conditions, in the context of on-farm Dynamic Management (OFDM) systems, crop populations are submitted to both natural selection and human-mediated selective pressures through different farmer practices (Enjalbert *et al.*, 2011). Several studies have shown that OFDM makes it possible to maintain a high level of diversity which is influenced by farmers’ uses and practices (Elias *et al.*, 2001; Pressoir and Berthaud, 2003a). One particular practice is seed exchange mediated by social organization that strongly reshapes the crop genetic diversity (Thomas *et al.*, 2012). Most of seed sources usually come from the same community and long-distance seed exchanges rarely occur (Louette *et al.*, 1997; Bellon *et al.*, 2011; Thomas *et al.*, 2011; Samberg *et al.*, 2013). Another common farmer’s practice is mixing seeds of several varieties and then resowing the mixture of the harvested seeds. Such mixtures of varieties are relatively widespread due to their year in, year out robustness that enables them to tolerate variations in biotic and abiotic pressures (Dawson and Goldringer, 2012). In a mixture of genotypes, adaptation can result from the increase in frequency of the most adapted component or from the emergence through recombination of a new genotype with higher fitness depending on the mating system. Note that in a mixture individual fitness would depend both on local adaptation and competitive ability. Yet, in a mixture of landraces, i.e. populations genetically heterogeneous, selection might occur within and among components and makes it even more complex. But up to now, little attention has been paid on the genetic mechanisms that underlie the micro-evolution of such populations simultaneously submitted to seed exchange, mixture and natural and human selection. Louette *et al.* (1997) suggested that crop populations under OFDM can be modelled as a metapopulation (Levins, 1969; Olivieri *et al.*, 1995). This model of crop metapopulation was further elaborated by adapting the general metapopulation model to the specific features of OFDM (van Heer-waarden *et al.*, 2010) and recently refined by Artoisenet and Minsart (2014). However in these two theoretical approaches, they chose to simplify the seed exchange process with a single parameter *m* accounting for all types of gene flows. Here, we alternatively considered that each seed diffusion event corresponds to a founding effect in order to better account for the underlying social organization. Such crop metapopulations are simultaneously submitted to natural selection and artificial selection operated by farmers, in addition to genetic drift and migration.

In this paper, we studied a recent mixture called ”Mélange de Touselles” (MDT) composed of three landraces of bread wheat (*Triticum aestivum*) and one landrace of cone wheat (*Triticum turgidum subsp. turgidum*). This mixture was created in 2001 by one farmer (HEF) and continuously cultivated on his farm since then. It was distributed in 2004 and in years after by HEF to other farmers. Thus, a set of MDT populations has been evolving within a group of farmers located in different sites in France, but also in Italy and in The Netherlands. To our knowledge, this original farmer-led design can be considered as the first on-farm evolutionary experiment.

In this context of evolutionary experiment, the main goal was to characterize the genetic structure and the spacial and temporal differentiation pattern of a self-pollinated crop mixture recently introduced in different environments and to identify some of the underlying evolutionary mechanisms. The evolutionary dynamics of the mixture was studied using on polymorphisms at neutral markers and at genes associated with flowering time variation. We focused on earliness as it is an important adaptive trait involved in the synchronisation of the plant cycle with the environment. A combine approach relying on a discriminant analysis and a dedicated network-based method was mobilized to decipher the complex genetic structure of the mixture.

A temporal and spatial sampling of MDT populations associated to genetic and phenotypic analysis allowed to i) describe the general genetic structure of the landraces composing the mixture, ii) estimate the differentiation of the mixture at the phenotypic, neutral genetic and gene levels, iii) understand the genetic and evolutionary mechanisms that underlie the differentiation and response to selection of the different populations. At last, this work contributed to over understanding of the combine role of farmers practices and of environment on the rapid evolution of a crop mixture and highlighted the crucial role played by the within-population variability in adaptation.

## 3 Materials and methods

### 3.1 Description of the on-farm evolutionary experiment

The mixture studied was developed by a farmer (HEF), who decided at the end of the 1990’s to re-establish four local landraces of the southeastern region of France. Landraces historically grown in that region disappeared in the middle of the twentieth century when modern agriculture replaced landraces and old varieties by elite material. He obtained from the INRA Clermont-Ferrand genebank around 50 seeds of each of the four varieties: *Touselle Anone* (TAN), *Touselle Blanche Barbue* (TBB), *Touselle Blanche de Provence* (TBP) and *Touselle sans Barbe* (TSB). It is important to note that TBB is an allotetraploid variety of wheat (*Triticum turgidum subsp. turgidum*) whereas the others are allohexaploid (*Triticum aestivum*). The landraces have been grown in small plots from 1997 to 2001, which allowed to increase plot size from 1 to 10, to 100 and to 1000 *m*^2^, respectively. In 2001, bad weather conditions caused important lodging and HEF decided to harvest the four varieties together. This mixture has been maintained since then in a large plot of around one hectare. In 2004, he started to distribute seed lots of the mixture to other farmers. We studied temporal samples of HEF as well as samples of mixture from 13 other farmers who had been given the MDT in 2004, 2005 and 2006 (see Table 1). Further spreading of the MDT mixture will not be considered in this study. This collective experience led by farmers themselves was thus studied as a case of on-farm evolution of a crop metapopulation.

**Table 1:** Description of the MDT samples (#G: Number of generations since the MDT was created by HEF, NBGENOUT: number of generations grown outside HEF’s farm).

| Sample name | Owner | Harvest<br>year | Farming practices | #G | NB<br>GEN<br>OUT | Long. | Lat. | Alt. (m) |
| --- | --- | --- | --- | --- | --- | --- | --- | --- |
| HEF03fld | HEF | 2003 | field | 2 | 0 | 4.340 | 43.790 | 109 |
| HEF05fld | HEF | 2005 | field | 4 | 0 | 4.340 | 43.790 | 109 |
| HEF06fld | HEF | 2006 | field | 5 | 0 | 4.340 | 43.790 | 109 |
| HEF07fld | HEF | 2007 | field | 6 | 0 | 4.340 | 43.790 | 109 |
| HEF08fld | HEF | 2008 | field | 7 | 0 | 4.340 | 43.790 | 109 |
| MIR08fld | MIR | 2008 | field | 7 | 4 | 4.030 | 44.020 | 131 |
| CER08mul | CER | 2008 | medium-sized plot | 7 | 4 | 5.370 | 44.900 | 1068 |
| MAI06fld | MAI | 2006 | field | 7 | 2 | -0.230 | 47.460 | 21 |
| MAI08fld | MAI | 2008 | field | 7 | 4 | -0.230 | 47.460 | 21 |
| CHD08fld | CHD | 2008 | field | 7 | 3 | 5.400 | 45.530 | 495 |
| RAB07col | RAB | 2007 | small plot | 6 | 2 | 6.310 | 46.170 | 500 |
| VIC08mul | VIC | 2008 | medium-sized plot | 7 | 3 | -1.130 | 47.010 | 90 |
| AUD08col | AUD | 2008 | small plot | 7 | 3 | 0.650 | 46.670 | 33 |
| BER08col | BER | 2008 | small plot | 7 | 3 | 5.270 | 47.560 | 296 |
| BER08mul | BER | 2008 | medium-sized plot | 7 | 3 | 5.270 | 47.560 | 296 |
| INE08col | INE | 2008 | small plot | 7 | 2 | 11.430 | 45.620 | 260 |
| OLR08col | OLR | 2008 | small plot | 7 | 2 | 1.710 | 49.120 | 127 |
| HZE08col | HZE | 2008 | small plot | 7 | 2 | 5.230 | 52.180 | -3 |
| ICE08col | ICE | 2008 | small plot | 7 | 2 | 13.900 | 42.020 | 577 |
| FLM07col | FLM | 2007 | small plot | 6 | 1 | -0.650 | 47.420 | 52 |
| FLM08col | FLM | 2008 | small plot | 7 | 2 | -0.650 | 47.420 | 52 |
| TANCLM04col | CLM | 2004 | small plot | - | - | - | - | - |
| TBBCLM03col | CLM | 2003 | small plot | - | - | - | - | - |
| TBPCLM04col | CLM | 2004 | small plot | - | - | - | - | - |
| TSBCLM03col | CLM | 2003 | small plot | - | - | - | - | - |

### 3.2 Data collection

Information about the diffusion of MDT was obtained through several interviews with HEF. Directed telephone interviews were carried out with 10 French farmers who grew the mixture in order to collect information about farming practices on the different farms. Similar information about the two populations grown in the Netherlands and the other one in Italy has been obtained from the Farm Seed Opportunity project in which this mixture had been distributed to Dutch and Italian farmers (Serpolay *et al.*, 2011). Based on this information, we built a seed diffusion and multiplication network (Figure 1) and we estimated demographic population size for each population. In addition, the genebank curator in charge of the wheat collection from INRA Clermont-Ferrand genebank (CLM) was also interviewed to gather information about multiplication and growing practices during seed multiplication and regeneration of the accessions. Specific information about management practices of TAN, TBB, TBP and TSB was also collected.

**Figure 1:**
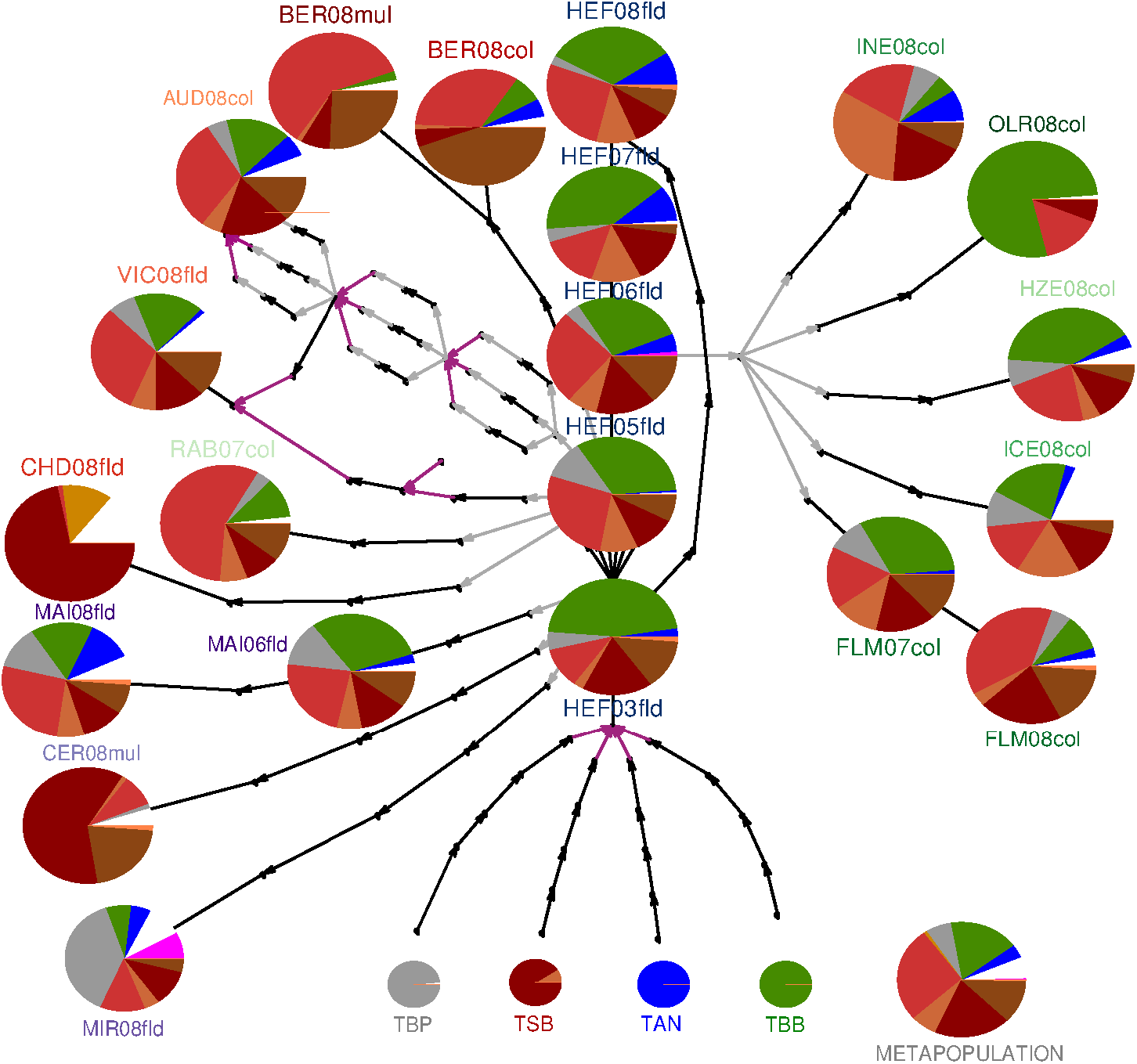
Frequency of the different genetic groups among the sampled populations. At the bottom, the genetic composition of the four reference landraces (TBP, TSB, TAN and TBB). The five vertically aligned pies are time series of HEF’s population. The other pies correspond to populations grown by different farmers for one to four generations. TSP is in gray, TSB is divided into five sub-groups (TSB1: red, TSB2: light red, TSB3: dark red, TSB4: brown and TSB5: salmon-pink), TAN is in blue, TBB in green, FLA in pink, THA in orange and UH in white. Multiplication events are represented by black arrows, diffusion by gray arrows and mixture by purple arrows. Pies size is proportional to the number of individuals: 32 plants were studied for the four reference landraces and almost 80 plants per populations.

#### Sampling strategy

To depict the evolution of the recent MDT mixture, 21 samples were collected from 14 farmers including HEF (Figure S1 in Supporting information). The name of the samples contains three types of information: (i) the name of the owner (first three characters); (ii) the harvest year; (iii) the farming practice (last three characters) (See Table 1 for more details). Thus, a MDT population is defined by the farm where it has been cultivated and the farming practices, *i.e.* information (i) and (iii). The material studied included five samples corresponding to a time series of HEF’s population (HEF03fld, HEF05fld, HEF06fld, HEF07fld, HEF08fld, respectively), two temporal samples for MAI’s population (MAI06fld, MAI08fld) and for FLM’s population (FLM07col, FLM08col). All other samples correspond to populations from different farms where the mixture has been grown for a time period ranging from one to four years. A snapshot of the MDT metapopulation was studied considering the subset of fifteen populations sampled in 2007 and 2008: HEF08fld, MIR08fld, CER08mul, MAI08fld, CHD08fld, RAB07col, VIC08fld, AUD08col, BER08col, BER08mul, INE08col, OLR08col, HZE08col, ICE08col, FLM08col. In addition, samples of the four varieties that were used to build the mixture (TAN, TBB, TBP and TSB) were provided by the INRA Clermont-Ferrand genebank (CLM) in order to have a reference for the initial composition of the MDT mixture.

#### Molecular analyses

In December 2008, seeds from the 21 MDT samples and from the four varieties from CLM were sown in pots in the greenhouse at Le Moulon experimental station. In January 2009 leaf samples were taken from 80 plants per population, while only 32 individuals were sampled for each of the four varieties, *i.e.* 1808 individuals. For each plant, total DNA was extracted from 100 mg of fresh material following a protocol derived from the QIAGEN DNeasy 96 Plant Kit (QIAGEN, Basel, Switzerland). Fourteen microsatellite markers (Single Sequence of tandem Repeats (SSRs)) developed by Röder *et al.* (1998), one (wmc231) by Somers *et al.* (2004) and a bi-locus marker (cfd71) developed by Guyomarc’h *et al.* (2002) were used for genotyping the 1808 individuals studied. This set of 17 markers covered 17 of the 21 chromosomes of bread wheat. Only chromosomes 1A, 6A, 6B and 7D were not covered. PCR protocols were adapted from Röder *et al.* (1998) and Guyomarc’h *et al.* (2002). Amplified fragments were separated on an ABI 3130xl semi-automatic sequencer (Applied Biosystems) and analysed with GeneMapper 3.7 (Applied Biosystems). Individuals with more than 6 missing data among the 17 SSR markers were discarded. After genotyping, the final dataset contained 1793 individuals out of the 1808 initial set. Sometimes bands were detected for markers mapped on the D genome in individuals from the initial TBB sample (allotetraploid genome AB) although SSR markers were designed to be specific for each genome (A, B and D, respectively). To avoid overestimation of interspecific crosses, bands that were specific of the initial TBB landrace were noted among the 17 markers to be used as a reference. Then, a screening was performed to detect and to separate TBB individuals from the whole dataset based on these signature bands. These TBB individuals were scored with a specific allele code for loci mapped in the D genome and were studied independently in some analyses.

Four polymorphisms located in candidate genes previously shown to be associated with earliness (Rousset *et al.*, 2011) were genotyped in order to detect if these genes were submitted to selection during the evolution of the mixture. One Single Nucleotide Polymorphism (SNP; C/T) located in the first intron of the FTA gene (Bonnin *et al.*, 2008) was genotyped, as well as three polymorphisms located in two of the three copies of the *VRN-1* gene (A, and D genomes):

- one polymorphism by duplication, insertion and deletion in the promoter region of *VRN-1A* (named *VRN-1A_prom_*), identified by Yan *et al.* (2004),
- one substitution in the seventh exon of *VRN-1A* (*VRN-1A*_*ex*7_), detected by Sherman *et al.* (2004),
- a four kb deletion in the first intron of *VRN-1D* (*VRN-1D*), detected by Fu *et al.* (2005)

#### Greenhouse experiment

In February 2009, plants were transplanted to ground soil under a plastic tunnel to be submitted to natural vernalization at the four leaf stage following a completely randomized two-block design. Viability, plant size and heading date as a proxy for flowering time were scored during plant development at the individual level. An individual was considered as having headed when half of the ear was out of the leaf sheath. It is important to note that 25% of the plants was quite tall in the experimental conditions. This might have led to competition among neighbouring plants.

### 3.3 Data analysis

#### Detection of the genetic structure

As wheat is highly autogamous, all the studied loci were in strong linkage disequilibrium (data not shown). For this reason, the fine population structure was studied considering each multilocus genotype as two haplotypes. Haplotype reconstruction and inference of missing data were performed on the 1793 individuals using PHASE software (Stephens *et al.*, 2001) using simultaneously the 17 SSR markers and the four polymorphisms of candidate genes. As suggested in (Garrick *et al.*, 2010), the model with recombination (MR) was used. The analysis was performed with 100 burn-in periods before 100 iterations and a recombination rate between loci equal to Then, pairs of haplotypes with the highest probability for each individual were selected. Focusing on the 17 SSR markers, this new dataset was called the phased Multi-Locus Genotype (pMLG). Among-and within-populations haplotype variation was calculated using Arlequin software (Excoffier and Lischer, 2010) by estimating the unbiased haplotype diversity (*H*_*D*_), which accounts for small population sizes (Nei, 1987).

Two methods were used to detect the structure of the genetic diversity based on the pMLG dataset: i) a Discriminant Analysis on Principal Component (DAPC) (Jombart *et al.*, 2010), ii) a haplotype network analysis specifically adapted for the purpose of the study. TBB individuals, which had been already identified due to their high level of divergence compared to the other varieties, were not included in the analysis. In addition, a few very similar individuals showed a pattern clearly different from the rest. They were identified as Florence-Aurore (FLA), an old variety also grown by HEF. It was detected using data at 13 out of the 17 SSR markers that have also been used in a previous diversity study where FLA was genotyped (Roussel *et al.*, 2004). These individuals were also discarded from the analysis.

DAPC was run using *adegenet* (Jombart, 2008), a package developed in R (R Core Team, 2014). DAPC was applied on the results of a k-means clustering (Hartigan and Wong, 1979). K-means is an unsupervised classification procedure that aims to partition *n* observations into *k* clusters in which each observation belongs to the cluster with the nearest mean. K-means was performed on data obtained after a Principal Component Analysis (PCA) on centered data. Different numbers of initial seeds and of iterations were tested for a range of group numbers (from k=2 to 10) with 10 replications each time. The k-means algorithm was the most convergent one for 160 initial seeds and 100,000 iterations for each value of k. The k value was chosen using the Bayesian Information Criterion (BIC). Then, the discriminant analysis was carried out on the principal components using the optimal number of groups detected previously.

The haplotype network analysis, an adaptation of the Rozenfeld’s method (Rozenfeld *et al.*, 2008), was performed using the number of differences among all pairs of unique haplotypes present in the dataset (Thomas *et al.*, 2012). This information was stored in a matrix (*A*) where *A*_*ij*_ is the number of differences between haplotypes *i* and *j*. Undirected networks were plotted where each node corresponded to a distinct haplotype and edges linked two haplotypes *i* and *j* only if *A*_*ij*_ ≤ *th*, with *th* the maximum number of differences between two haplotypes, *th* varying from 1 to 17. Networks were drawn with the Pajek software (Batagelj and Mrvar, 2002). Kamada-Kawai’s force-based algorithm (Kamada and Kawai, 1989) was used to define the spatial distribution of the nodes. The threshold was set to two (*B*^(2)^) so as to detect haplotypes specific of each of the initial four varieties (TAN, TBB, TBP and TSB) as independent genetic groups. Then, each haplotype non-specific of the four initial varieties but connected to one of the four genetic groups was assigned to this group. A new genetic group was defined for every set composed of a minimum of five distinct haplotypes. Other haplotypes were defined as unassigned haplotypes (UH). The analyses have been implemented in R (R Core Team, 2014) and the scripts are available on request.

The results of the two clustering methods (DAPC and the haplotype network analysis) were compared and summarized in the form of pseudo-alleles corresponding to the different genetic groups and called virtual multi-allelic marker (VMK). In order to connect genetic data and phenotypes, haplotype information was assembled into genotypes. For that, we assigned each individual to a genotype group (GENOGP) resulting from the two pseudo-alleles. The genotype group (GENOGP) of each individual was defined based on its two pseudo-alleles. For instance, if the genetic group TBB was detected twice in the same individual, then its genotype group was set to TBBTBB. Only heterozygotes falling into two distinct genetic groups were defined as heterozygous at this marker. The VMK was used to follow the evolution of the composition of each population in terms of group frequencies.

#### Within and among population diversity

Genetic diversity of the 1793 pMLG distributed in the 21 MDT samples and of the four initial varieties was studied at the level of the 17 SSR markers, at the VMK level defined in the previous section and at the level of the four polymorphisms in candidate genes (*FTA*, *VRN-1A_prom_*, *VRN-1A*_*ex*7_ and *VRN-1D*). Unbiased Nei’s estimate of genetic diversity (*H*_*E*_) (Nei, 1978), mean observed heterozygosity (*H*_*O*_), allele richness (*R*_*S*_) and the deviation from Hardy-Weinberg genotypic proportions (*F*_*IS*_) were estimated with Genetix software (Belkhir *et al.*, 2000). Haplotype richness (*HR*) was computed as the number of unique haplotypes in a population divided by twice the number of individuals in the population. Haplotype diversity (*H*_*D*_) was computed as the equivalent of *H*_*E*_ for haplotypes (Nei, 1987).

Differentiation among populations was calculated using *θ*, the *F*_*ST*_ estimator developed in Weir and Cockerham (1984) and implemented in the Genetix software (Belkhir *et al.*, 2000).

#### Effective population size estimation

The genetic effective population size (*N*_*E*_) was estimated using the temporal variation of allele frequencies for each of the 17 SSRs (Waples, 1989). *N*_*E*_ is given by:

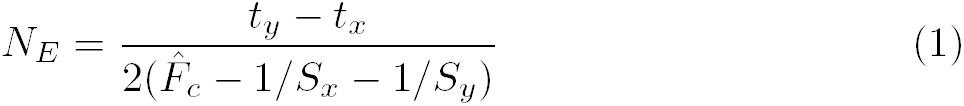

where *S*_*x*_ is the number of individuals sampled at the *t*_*x*_ generation (respectively *S*_*y*_ individuals at *t*_*y*_) and 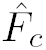 is the variance in allele frequency defined by:

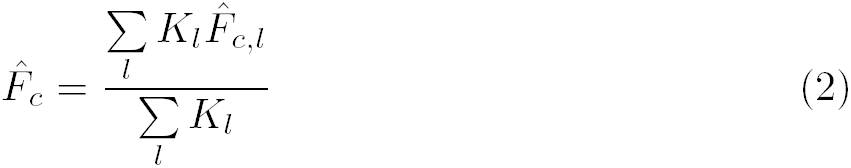

where *K*_*l*_ is the number of alleles at locus *l* and 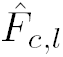:

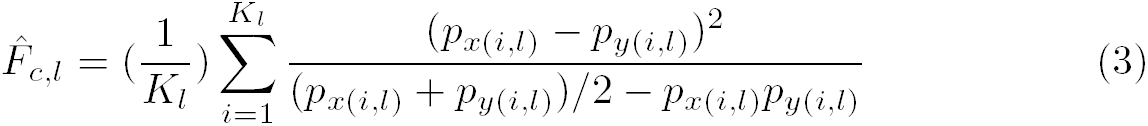

where *p*_*x*(*i,l*)_ [respectively *p*_*y*(*i,l*)_] represents the frequency of allele *i* at locus *l* in the sample of *S*_*x*_ individuals drawn at generation *t*_*x*_ (respectively *S*_*y*_ individuals at *t*_*y*_).

#### Statistical analyses of earliness

Fisher’s exact tests on the number of viable plants and headed plants were performed using R (R Core Team, 2014). SAS/STAT software, Version 9.2 of the SAS System for Unix (Copyright © 2002-2008 SAS Institute Inc) was used for the other statistical analysis. The analyses of covariance (ANCOVA) were performed using the Generalized Linear Model (GLM) procedure. The test for the normal distribution of the residuals was tested using the UNIVARIATE procedure and the variances were estimated using the VARCOMP procedure with the restricted maximum likelihood (REML) approach.

The basic initial model addressed the within-trial micro-environmental variations due to the experimental conditions. To account for potential competition among neighboring plants, a neighborhood covariate (NBH) was computed for each plant as the difference between the mean plant height of the eight neighboring plants and the plant height of the considered individual plant.

The effect of the replicated block (*REP*) and of neighborhood (*N BH*) were tested as follows:

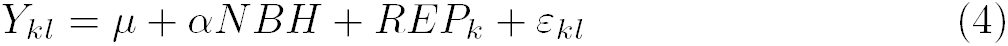

where *Y*_*kl*_ is the heading date value of plant *l* in *REP k*, *N BH* is a continuous variable, *REP* the fixed block effect, and *ε_kl_* the random residual variable 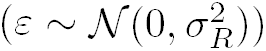.

The environmental variance 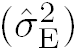 of the trial was estimated as the residual variance of the following model:

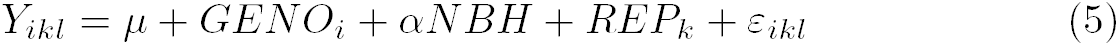

where the random *GENO* (*i* = 1..11) effect corresponds to the eleven most frequent homozygous genotypes (*GENO*) in the whole dataset. Thus, the residual variance of this model corresponded to between-plant environmental variability.

#### Spatial differentiation

We then tested for differentiation among populations within the MDT metapopulation using the following model:

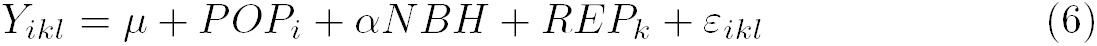

where *POP* (*i* = 1*…*15) is the random population effect corresponding to the 15 MDT populations sampled in 2007 and 2008.

In this model, the estimated variance of the POP effect gave the among-population genetic variance 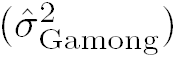. As the residual variance of the model included both within-population genetic variance 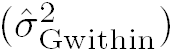 and among-plants environmental variation 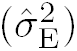,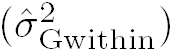 was obtained as follows:

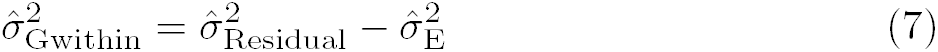

with 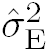, the environmental variance estimated in model (5).

To compare differentiation at a quantitative trait level to differentiation at the neutral marker level (*F*_*ST*_), *Q*_*ST*_ (Wright, 1969; Spitze, 1993) was estimated for the metapopulation subset:

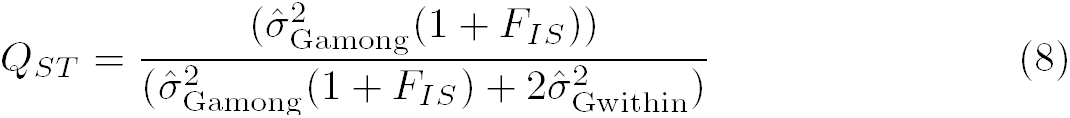

As wheat is mainly a selfing species, we assumed that *F*_*IS*_ ⋍ 1, and we obtained:

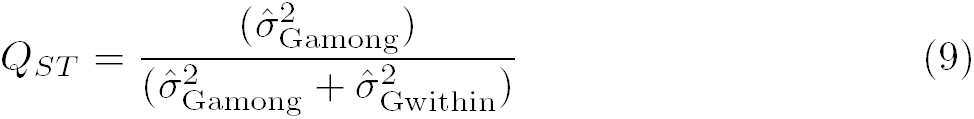

A multiple regression model was used to assess whether the average heading date of the different populations could be explained by environmental characteristics of the cultivation sites such as latitude (*LAT*), longitude (*LONG*), altitude (*ALT*), population size (*SIZE*), and the number of generations during which the mixture had been grown on a given farm after moving from HEF’s (*NBGENOUT*). The multiple regression was performed using a stepwise model with the REG procedure and the FORWARD method.

The role of genetic structure in earliness differentiation was investigated at the population level. A regression analysis was carried out to test whether the frequency of the genotype groups composing the samples explained the average heading date.

The effect of genetic structure was also investigated at the individual level to test whether phenotypic differences were due to the genotype groups:

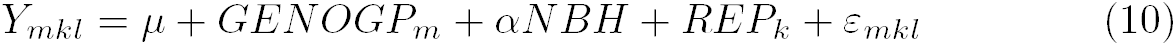

with *GENOGP* a random effect (*m* in 1…7), characterizing each of the main genotype groups detected in section *Detection of the genetic structure* (DAPC and haplotype network analysis) to limit the number of unbalanced classes.

Then, model (6) was run on data subsets composed of each main genotype group in order to estimate the among-population within-groups genetic variance 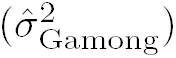 for each of them and to compare it with their genetic diversity (*H*_*E*_).

#### Association study between heading date and candidate genes for earliness

The effects of candidate gene polymorphisms (*M K_j_*) on heading date were assessed at three levels, considering :

- the population effect:

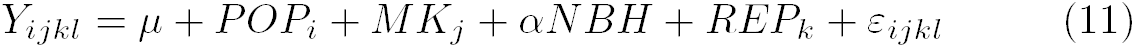
- the genetic background (GENOGP effect):

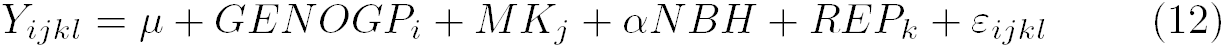
- the population effect within a given genotype group: model (11) was run for each of the main genotype group.

Each polymorphism (*M K*: *VRN-1A_prom_*, *VRN-1D*, *VRN-1A_ex7_* and *FTA*) was tested both separately and together.

## 4 Results

### 4.1 Genetic structure of the MDT populations

#### Genetic diversity : initial *vs* metapopulation level

The evolution of the Mélange de Touselles diversity was studied by comparing:

- the four reference varieties (TAN, TBB, TBP and TSB) pooled together with equal frequency, named virtual MDT since the initial frequencies of the four components were not known (41 distinct haplotypes)
- the initial reference population, first mixture available, HEF03fld (62 distinct haplotypes),
- the MDT metapopulation composed of the fifteen populations sampled in 2007 or 2008 (440 distinct haplotypes).

Virtual MDT showed a slightly lower diversity at neutral markers (expected heterozygocity, *H*_*E*_ = 0.49 and haplotype diversity, *H*_*D*_ = 0.88) as compared to HEF03fld (*H*_*E*_ = 0.51 and *H*_*D*_ = 0.94, Table S1 in Supporting Information). The number of shared haplotypes between the virtual MDT and HEF03fld was quite low (only seven distinct haplotypes) with 82.9% of the haplotypes initially present in the virtual MDT not detected in the HEF03fld. Reversely, the HEF03fld population was composed of 88.7% of haplotypes not present in the virtual MDT, among which 61.3% were recombinant haplotypes among haplotypes present in the virtual MDT and 27.4% were new haplotypes with one to three new alleles per individuals undetected in the virtual MDT (Figure 2A.). Consistently, a highly significant pairwise *F*_*ST*_ was estimated between Virtual MDT and HEF03fld (*F*_*ST*_ = 0.56).

**Figure 2:**
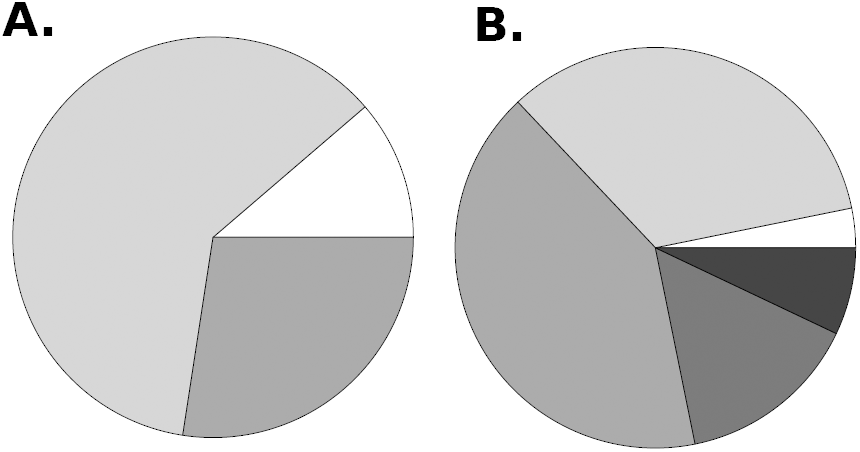
Distribution of the unique haplotypes observed in : A. HEF03fld (62), B. MDT metapopulation (440). Shared haplotypes with Virtual MDT are in white, recombinant haplotypes of Virtual MDT are in light grey, haplotypes with new allele(s) are in medium grey, shared haplotypes with HEF03fld are in dark grey, recombinant haplotypes of HEF03fld are in black.

Genetic diversity at the metapopulation level (*H*_*E*_ = 0.51) was similar to the initial diversity in HEF03fld (*H*_*E*_ = 0.51) (Table S1). The metapopulation showed a slightly higher haplotype diversity compared to HEF03fld (*H*_*D*_ = 0.98), and a higher allelic richness (*R*_*S*_ = 9.35 *versus R_S_* = 3.65 for HEF03fld). Only fourteen distinct haplotypes among the 440 detected in the metapopulation were shared with the virtual MDT. Within the 97% of haplotypes of the metapopulation not present in virtual MDT, 7% were shared with HEF03fld, 34% were recombinants of virtual MDT, 15% were recombinants of HEF03fld and 41% were haplotypes with new alleles neither present in Virtual MDT nor in HEF03fld (Figure 2B.). A lower but significant pairwise *F*_*ST*_ was estimated between HEF03fld and the metapopulation (*F*_*ST*_ =0.009). The genetic parameters of the MDT populations revealed two groups of populations. Five populations (CER08mul, BER08col, BER08mul, CHD08fld, OLR08col) had a lower diversity than HEF03fld and metapopulation for *H*_*E*_ and *H*_*D*_, while the other ones had the same level of diversity than HEF populations (Table S1).

#### Genetic structure of the MDT populations

A clustering analysis on the 1793 pooled pMLG was performed at the haplotype level working with 517 unique haplotypes. The samples of the four landraces provided by the genebank were initially used as references to define the boundaries of each variety. Because TBB is a tetraploid wheat species, TBB related individuals were easily identified and isolated from the rest of individuals. It was more difficult for the rest since, as previously shown, most of the haplotypes observed in the dataset were not present in the four reference samples. Moreover, none of the four landraces was completely homogeneous and the two most heterogeneous TBP and TSB were quite close genetically, causing a lack of discrimination. We were thus faced with a tricky population structure with populations composed of different varieties that were in turn composed of several genetic groups with some overlaps among varieties.

Two complementary clustering methods namely DAPC and a haplotype network-based method, were used to disentangle the structure of the populations at the haplotype level. The clustering was performed after discarding the TBB (407 individuals) and FLA groups (7 individuals) previously identified. K-means algorithm was performed with k, the number of clusters between two and 20 groups. Six groups (k=6) showed the minimal BIC value with a strong and stable elbow compared to the other k values. Based on this clustering, DAPC provided the probability of assignation of each individual to each of the k groups. Two genetic groups were easily assigned to TAN and TBP on the basis of the reference varieties. The sample of the TSB reference variety for TSB was subdivided into two groups (further named TSB2 and TSB3). At this stage, two other groups were still not assigned.

The haplotype network also allowed to detect TAN and TBP (Figure 3) but for TSB only one main group and a closely related minor one (named TSB5) were detected. A new subgroup was detected within the TSB3 group. It was only observed in the CHD08fld population. Interviews revealed that in 2006 this population was exposed to a mixture with another landrace called *Touselle des Hautes-Alpes*. Therefore this TSB group was assumed to be derived from this landrace and was called THA even though no sample was available to check if we actually detected that particular variety. The remaining TSB3 individuals were maintained as the TSB3 group. Combining both DAPC and haplotype network approaches led to the conclusion that the two unknown groups detected with DAPC most likely fell into the TSB variety. They will further be named TSB1 and TSB4.

**Figure 3:**
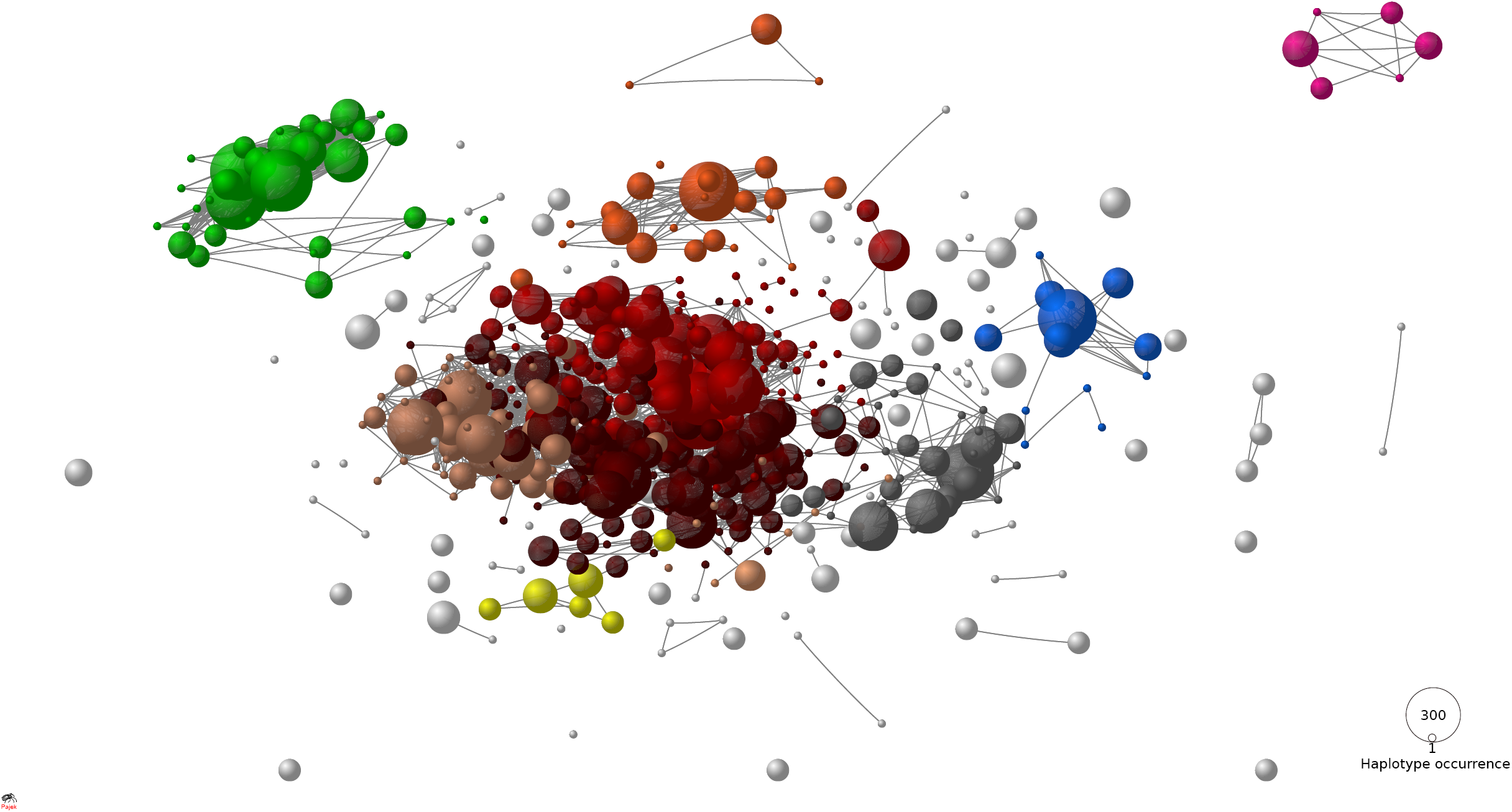
Global haplotype network composed of 517 distinct haplotypes (nodes). Two haplotypes are connected if they show two or less differences. Spatial distribution of the nodes accounts for the total number of differences between each pair of haplotypes. Each color corresponds to a different genetic group. TSP is in grey, TSB is divided into five sub-groups (TSB1: orange, TSB2: red, TSB3: dark brown, TSB4: light brown and TSB5: salmon-pink), TAN is in blue, TBB in green, FLA in pink, THA in yellow and UH in white. FLA and THA are genetic groups not present in the initial mixture and UH corresponds to the unassigned haplotypes. Node size is proportional to the number of haplotypes found in the pooled dataset (1793 pMLG, i.e.: 3586 haplotypes).

Haplotypes belonging to the same genetic group (of the same color) appeared connected and close to each other while most of the unassigned haplotypes (UH) were on the outskirts of the network, consistently with the purpose of the method (Figure 3). Indeed, the haplotype network method allowed us to detect each haplotype which did not fall into one of the genetic groups. They were labelled as unassigned haplotypes (UH), acknowledging it is complex to decipher between recombinant, migrant or experimental artifacts. Further investigations would be needed to determine the nature of these haplotypes. In this paper, we focused on the haplotypes belonging to the 10 identified genetic groups: TAN, TBB, TBP, TSB1, TSB2, TSB3, TSB4, TSB5, THA, FLA. They represent 94.0% of the 2234 haplotypes sampled within the MDT metapopulation, corresponding to 96% of the 1793 individuals. Because the UH haplotypes were not considered in this study, the role of farmers practices on the migration process among populations within the same farm will not be analysed.

#### Diversity of the initial four landraces

We analyzed the genetic diversity of the initial four landraces (Table 2) in terms of the 8 genetic groups (TAN, TBB, TBP, TSB1, TSB2, TSB3, TSB4, TSB5). Each landrace belonged to one genetic group except TSB which was divided into two groups, TSB2 and TSB3. TAN, TBB and TBP presented lower values of genetic diversity compared to TSB whatever the index (*H*_*E*_ or *R*_*S*_) and the sample (Virtual MDT, HE03fld or the metapopulation, Table 2).

**Table 2:** Summary statistics obtained for the four landraces considering each haplotype as one homozygote individual (N: number of haplotypes, *H*_*E*_: unbiased expected heterozygocity, *R*_*S*_: average number of alleles per locus).

| Variety | Haplotype | Virtual MDT |  |  | HEF03fld |  |  | Metapopulation |  |  |
| --- | --- | --- | --- | --- | --- | --- | --- | --- | --- | --- |
| | Group | $N$ | $H_E$ | $R_S$ | $N$ | $H_E$ | $R_S$ | $N$ | $H_E$ | $R_S$ |
| TAN | = | 64 | 0.010 | 1.060 | 4 | 0.000 | 1.000 | 89 | 0.040 | 1.880 |
| TBB | = | 64 | 0.030 | 1.240 | 74 | 0.050 | 1.350 | 417 | 0.050 | 2.350 |
| TBP | = | 63 | 0.030 | 1.290 | 8 | 0.080 | 1.240 | 153 | 0.060 | 2.180 |
| TSB |  | 64 | 0.260 | 2.180 | 74 | 0.340 | 2.820 | 1543 | 0.340 | 5.470 |
| - | TSB1 | 0 | - | - | 18 | 0.170 | 1.590 | 634 | 0.160 | 3.350 |
| - | TSB2 | 6 | 0.050 | 1.120 | 4 | 0.060 | 1.120 | 158 | 0.040 | 2.470 |
| - | TSB3 | 58 | 0.180 | 1.710 | 28 | 0.210 | 1.760 | 457 | 0.220 | 3.940 |
| - | TSB4 | 0 | - | - | 22 | 0.100 | 1.470 | 286 | 0.110 | 2.590 |
| - | TSB5 | 0 | - | - | 2 | 0.040 | 1.060 | 8 | 0.120 | 1.350 |

#### Differences among genetic groups

Pairwise *F*_*ST*_ between genetic groups were very high, ranging from 0.47 to 0.94 (Table 3). TBB showed the highest level of divergence with other genetic groups, which is consistent with the fact that TBB belongs to a different wheat species (*Triticum turgidum*). When all TSB genetic groups were pooled together, this group showed the lowest level of differentiation compared to the other genetic groups probably due to the high within-landrace diversity. However, each of the five TSB groups highly diverged from other groups with *F*_*ST*_ = 0.61. Therefore, we chose to work at the genetic group level instead of the initial landrace level, since the genetic group seemed to be a relevant evolutionary unit to follow the diversification process of MDT.

**Table 3:** Pairwise *F*_*ST*_ s between the genetic groups of the whole dataset (*N* = 3717 inferred haplotypes): all the MDT populations and reference samples from CLM (VIRT MDT). The pairwise *F*_*ST*_ s were estimated based on the multilocus frequencies.

| Pairwise $F_{ST}$ | TAN | TBB | TBP | TSB | TSB1 | TSB2 | TSB3 | TSB4 | TSB5 |
| --- | --- | --- | --- | --- | --- | --- | --- | --- | --- |
| TAN | 0.000 | 0.938 | 0.838 | 0.561 | 0.766 | 0.911 | 0.727 | 0.882 | 0.942 |
| TBB | - | 0.000 | 0.894 | 0.697 | 0.859 | 0.932 | 0.846 | 0.915 | 0.924 |
| TBP | - | - | 0.000 | 0.472 | 0.667 | 0.851 | 0.612 | 0.801 | 0.762 |
| TSB | - | - | - | 0.000 | - | - | - | - | - |
| TSB1 | - | - | - | - | 0.000 | 0.735 | 0.451 | 0.582 | 0.602 |
| TSB2 | - | - | - | - | - | 0.000 | 0.694 | 0.834 | 0.885 |
| TSB3 | - | - | - | - | - | - | 0.000 | 0.578 | 0.510 |
| TSB4 | - | - | - | - | - | - | - | 0.000 | 0.780 |
| TSB5 | - | - | - | - | - | - | - | - | 0.000 |

### 4.2 Evolution of the genetic composition across the metapopulation

At the metapopulation scale, the proportions of the different groups were more balanced than in the reference population (HEF03). On average, TBB decreased in frequency while the TSB groups became more frequent, with a particular increase of TSB1. However, group composition varied drastically from one population to the other (Figure 1). Sixteen out of the 21 populations were composed of more than 50% of TSB groups. TAN was maintained at a low frequency in most of the populations except in CER08mul, CHD08fld, RAB07col, BER08mul and OLR08col where it was not detected. TBP was present at a low frequency in most of the populations except in the MIR08fld population where it was much higher (30%), and in CER08mul, CHD08fld, BER08col, BER08mul, OLR08col where it was not detected.

Two levels of spatial and temporal differentiation among population were investigated: pairwise *F*_*ST*_ were estimated based on multilocus data and pairwise *F*_*ST*_ estimated based on the VMK to account for variations in genetic group composition. Pure temporal evolution of MDT was only captured by the time series from the HEF’s population. The results indicated a slight but significant temporal differentiation of MDT after 5 generations in HEF’s farm (*F*_*ST*_ (multilocus) = 0.014 and *F*_*ST*_ (VMK) = 0.011). The other sampled populations corresponded to independent evolutions of the initial HEF03fld mixture. The differentiation estimated at the metapopulation level (*F*_*ST*_ (multilocus)=0.111, *F*_*ST*_ (VMK)=0.158) indicated a high and significant differentiation among the set of populations harvested in 2007-08.

#### 4.2.1. Genetic effective population size

The genetic effective population size (*N*_*E*_) was estimated based on the temporal variations in allelic frequencies between each sampled populations and HEF’s population at the previous generation (Table S2 in supporting information). The effective population size estimated HEF’s farm fluctuated from 10 to 30 individuals except between 2005 and 2006 when 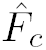 was of same order as the sampling effects. For this reason the average value was higher (*N*_*E*_ = 145). Almost all populations showed a *N*_*E*_ value of the same order of magnitude as HEF’s population ranging from 5 to 32, whatever the number of generations. ICE08col and AUD08col showed a slightly higher effective population size (*N*_*E*_ = 62 and *N*_*E*_ = 76 respectively). MAI06fld sample had the highest value (*N*_*E*_ = 153). It was not possible to estimate *N*_*E*_ for FLM07col, MAI08fld and VIC08fld because 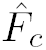 variations were too small compared to sample size. In this particular case, allele frequencies were stable after the diffusion event, unlike in the other populations.

### 4.3 Evolution of earliness

#### Phenotypic evolution

In order to connect genetic data and phenotypes, genotype groups instead of genetic groups will be considered in the following. As the experimental conditions were quite stressful for the plants, 25% did not reach heading. Fisher’s exact test was performed on the distribution of the number of viable plants and headed plants to detect if certain genotype groups or populations were more specifically affected by the growing conditions. The two tests were highly significant (*P*_*value*_ < 0.0001) at the genotype group (GENOGP) level as well as population (POP) level. The TBBTBB group was the most severely affected by these particular conditions (28.5% of the TBBTBB plants did not reach heading stage, to be compared with 25% overall).

The simple model (4), with a *REP* effect and the neighborhood covariate (*N BH*), explained 1.27% of the total variation of the heading date. REP effect was not significant whereas *N BH* covariate was highly significant (*P*_*value*_ < 0.0001) with a positive coefficient 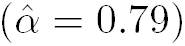 indicating that competition due to plant height differences among neighboring plants significantly delayed heading.

Metapopulation differentiation was studied with model 6 using all the pop*s*ulations sampled in 2008, plus RAB07col in 2007. The *POP* effect was highly significant (*R*^2^ = 12.7%, *P*_*value*_ < 0.0001), indicating among-populations divergence for earliness. MIR08fld, INE08col, ICE08col and BER08col populations were significantly earlier than OLR08col and RAB07col (with on average 6.4 to 9.1 days’ delay, Figure 4). The earliness of the other populations ranged between these extremes. 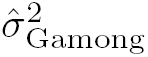 and 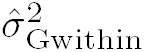 (equation (7)) were estimated to compute *Q*_*ST*_ (equation 9). The *Q*_*ST*_ value (0.26) was much higher than the multilocus *F*_*ST*_ value (0.11). This result indicated that differentiation was faster and larger for earliness than for neutral markers.

**Figure 4:**
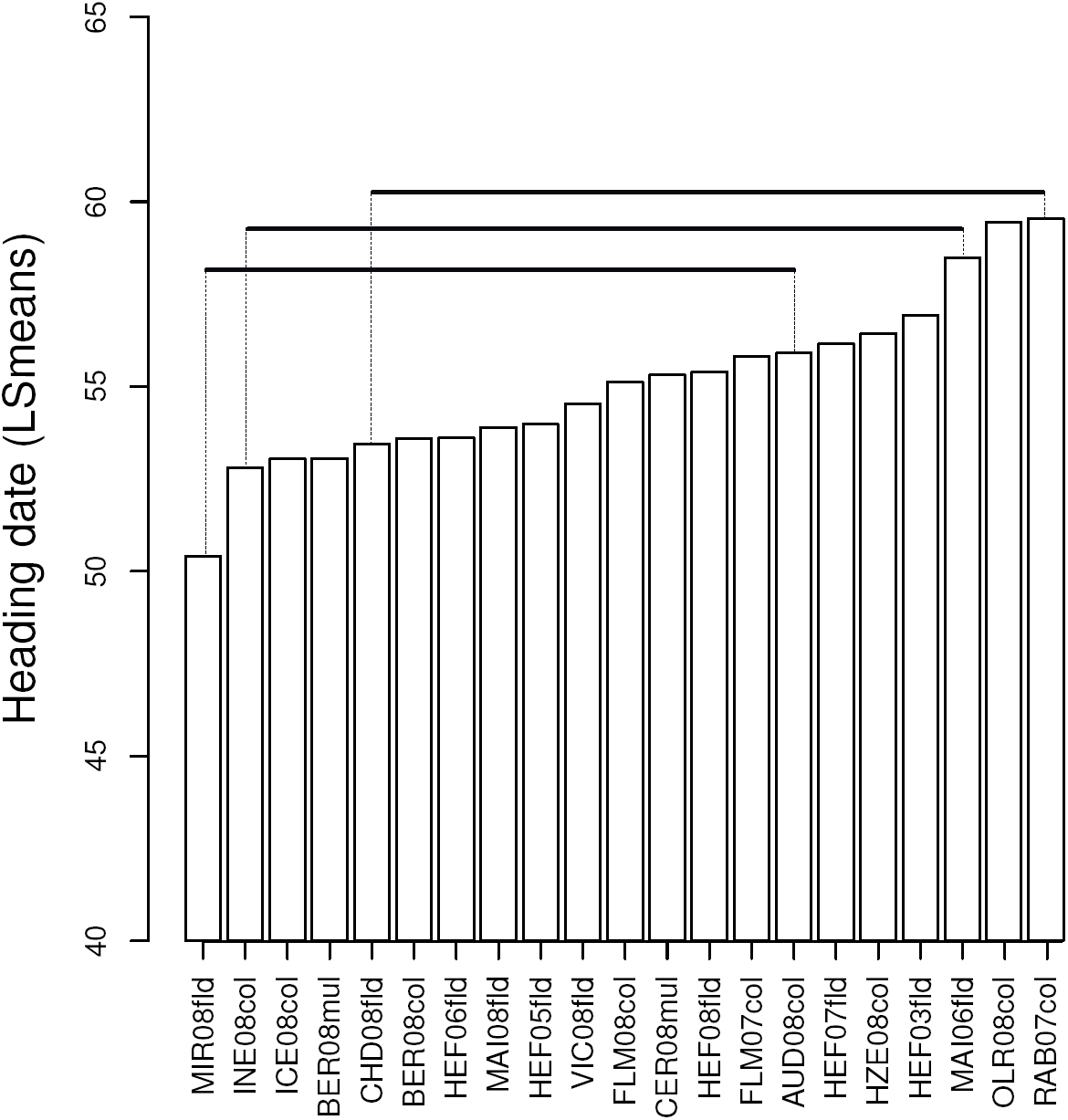
LSmean for heading date calculated for each population. The three lines above the bars indicate populations sharing the same statistical group

Five environmental continuous variables were tested in a step-wise multiple regression model to assess whether they explained the average heading date of the population. The best model explained 7.0% of the variability. The effect of demographic population size was not significant, while latitude, longitude, altitude, the number of generations outside HEF’s farm were significant. Latitude was the most significant parameter (partial *R*^2^ = 2.4%) with a positive value for the estimated coefficient. This indicated that populations located in the South became slightly earlier than populations grown in the North. Therefore, a large part of earliness among-population differentiation was due to other factors.

#### Genetic bases of heading dates differentiation

The aim of the following analyses was to assess the relative contribution of (i) variation in genotype group frequencies and of (ii) within group evolution to the population differentiation detected for earliness.

In the regression of the mean heading date of the populations on the frequency of each genotype group within population only TBBTBB had a significant positive effect (Figure 5). But in general, the mean heading dates of the populations were only weakly influenced by the frequencies of the genotype groups.

**Figure 5:**
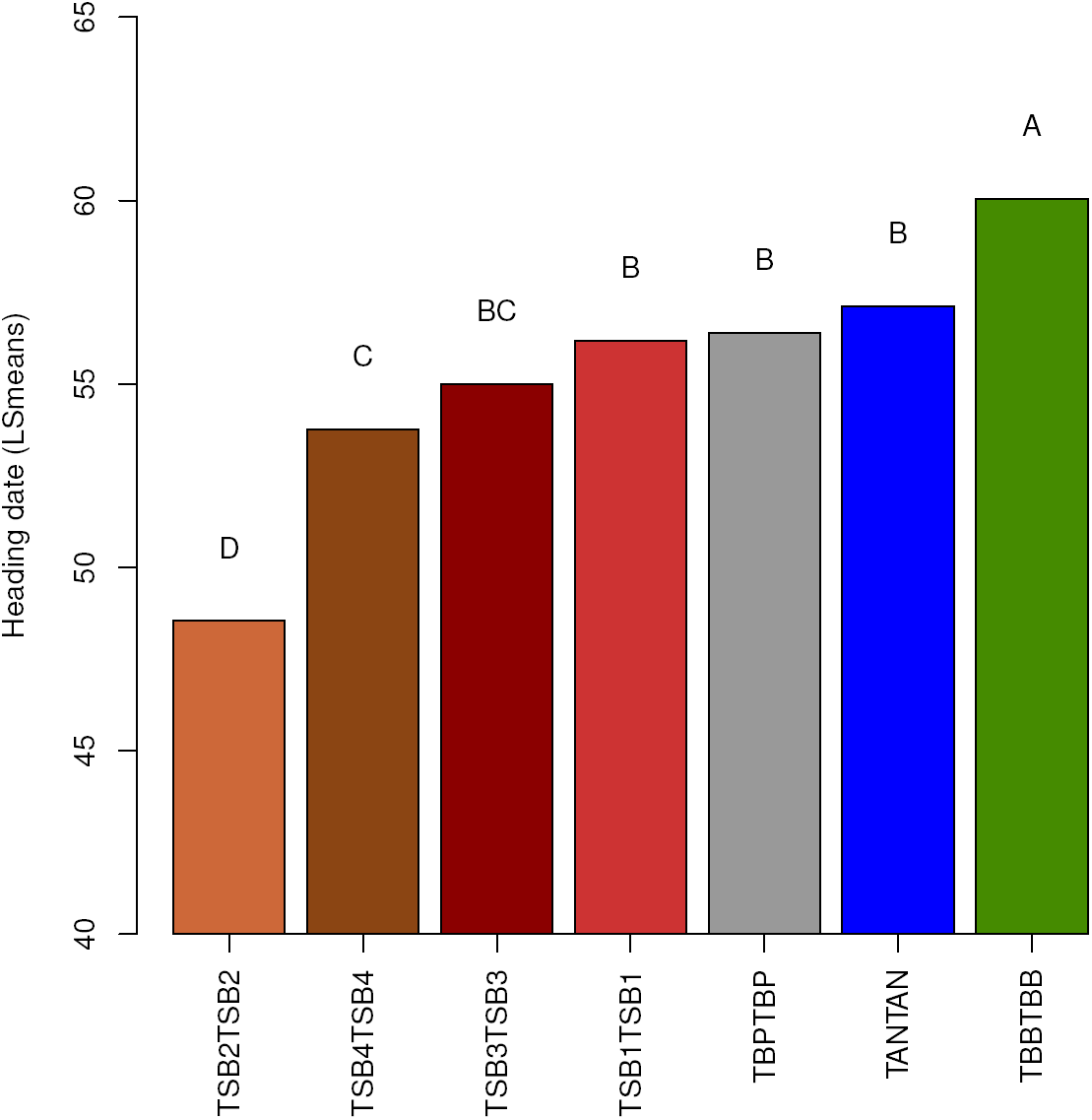
Weighted means for heading dates estimated for each genotype group. Significance of the tests was provided after Bonferroni’s multiple correction.

The effect of the genetic structure was tested on individual heading date using the genotype groups information. Here, only the seven most frequent genotype groups (TANTAN, TBBTBB, TBPTBP, TSB1TSB1, TSB2TSB2, TSB3TSB3 and TSB4TSB4) were considered (*GENOGP*). Note that this selection discarded heterozygotes that belonged to different genetic groups, but did not discard heterozygotes that belong to the same genetic group. Including *GENOGP* instead of *POP* in the analysis of covariance (equation 10 *versus* equation 6) dramatically increased the explanation in Table 4. The effect of *GENOGP* was highly significant (*P*_*value*_ < 0.0001), with substantial differences among weighted means (LSmeans) for genotype groups (Table 4 and Figure 5). TSB2TSB2 was the earliest genotype group (*HD* = 48.5) while TBBTBB was the latest (*HD* = 60.0), the others (TSBTSBs, TBPTBP and TANTAN) showing intermediate values.

**Table 4:** ANCOVA table for heading date considering the effects of *POPGP*, *POP*(*POPGP*) and *GENOGP*. *: *P*_*value*_ < 0.05, **: *P*_*value*_ < 0.01 and ***: *P*_*value*_ < 0.0001, -: not computed in the current model

| Model | *NBH* | | *REP* | | *POP* | | *GENOGP* | | $R^2$ |
| --- | --- | --- | --- | --- | --- | --- | --- | --- | --- |
|  | d.f. | F-value | d.f. | F-value | d.f. | F-value | d.f. | F-value |  |
| (6) | 1 | 15.05*** | 1 | 0.83 | 21 | 7.77*** | - | - | 0.12 |
| (10) | 1 | 54.63*** | 1 | 0.79 | - | - | 6 | 77.08*** | 0.30 |

Finally, we studied the among-population variation for each genotype group (model (6), Table 5). TSB2TSB2 showed the highest among-population genetic variance 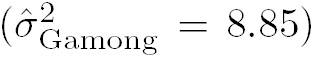 and TBPTBP the lowest 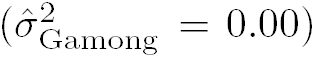 with a non-significant POP effect.

**Table 5:** ANCOVA table for heading date obtained with model (6) run for each genotype group and considering *POP* effects. *: *P*_*value*_ < 0.05, **: *P*_*value*_ < 0.01 and ***: *P*_*value*_ < 0.0001.

| Genotype group | *NBH* | | *REP* | | *POP* | | $\hat{\sigma}_{\text{Gamong}}^2$ | Residual<br>$\hat{\sigma}^2$ | $R^2$ |
| --- | --- | --- | --- | --- | --- | --- | --- | --- | --- |
|  | d.f. | F-value | d.f. | F-value | d.f. | F-value |  |  |  |
| TANTAN | 1 | 0.89 | 1 | 0.00 | 15 | 1.42 | 4.70 | 21.52 | 0.45 |
| TBBTBB | 1 | 18.57*** | 1 | 0.31 | 19 | 4.29*** | 8.31 | 18.75 | 0.32 |
| TBPTBP | 1 | 5.86* | 1 | 0.18 | 16 | 1.46 | 0.00 | 22.74 | 0.30 |
| TSB1TSB1 | 1 | 6.62* | 1 | 0.67 | 21 | 2.92*** | 2.89 | 25.41 | 0.19 |
| TSB2TSB2 | 1 | 6.93** | 1 | 0.88 | 19 | 2.78** | 8.85 | 25.18 | 0.38 |
| TSB3TSB3 | 1 | 3.12 | 1 | 0.08 | 21 | 1.58* | 1.22 | 20.81 | 0.15 |
| TSB4TSB4 | 1 | 3.20 | 1 | 0.34 | 19 | 2.65** | 7.56 | 23.07 | 0.32 |

#### Differentiation at earliness candidate genes

Four polymorphisms located in three earliness candidate genes (*FTA*, *VRN-1A* and *VRN-1D*) were studied. Genetic diversity was in general lower than at the neutral markers with similar trends over populations (Table S1).

Differentiation in the candidate genes associated to earliness (*F*_*STQ*_=0.239) was similar to differentiation for the quantitative trait (*Q*_*ST*_ =0.261) and much larger than neutral allelic differentiation (*F*_*ST*_ (multilocus)=0.111) and genotype groups differentiation (*F*_*ST*_ (virtual)=0.158). This trend might indicate that populations were submitted to divergent selection for earliness in relation to the contrasted environmental conditions or farming practices on the different farms. Phenotypic differentiation might be partly underlied by differentiation at *FTA*, *VRN-1A* and *VRN-1D*.

#### Association between heading date and candidate genes

The effect of each gene polymorphism was tested one by one as well as all together with the *POP* effect in equation (11). These models accounted for 16.0 to 36.5% of variability, with a strong significant *POP* effect. *FTA* and *VRN-1A*_*ex*7_ were not significant whereas *VRN-1D* and *VRN-1A_prom_* were highly significant when tested one by one (*P*_*value*_ < 0.0001, data not shown). These two genes were in general associated to the heading date whatever the population. Only *VRN-1A_prom_* remained significant when the four genes were tested all together. Model (12) allowed to assess the association between the earliness genes and the heading date while accounting for the genetic structure of the genotype groups. The models explained between 28.3 and 30.7% of the variability and *GENOGP* was always highly significant (*P*_*value*_ < 0.0001) (Table 6). The effects of *VRN-1A*_*ex*7_ and *VRN-1D* were not significant, whereas *FTA* and *VRN-1A_prom_* were significant (*P*_*value*_ < 0.05 and *P*_*value*_ < 0.0001, respectively). *FTA* marker was not associated in the previous model due to significant interactions between genotype groups and *FTA* alleles (data not shown). TSB1TSB1, TSB3TSB3 and TSB4TSB4 preferentially carried out the allele 1 of *FTA* whereas TANTAN, TBBTBB, TBPTBP and TSB2TSB2 carried out allele 2. Nevertheless, the same average value of heading date was observed for these two allele-specific groups. Model (11) was also tested in each genotype group. *FTA* was significant within TSB1TSB1 and TSB3TSB3 and *VRN-1A_prom_* was significant within TSB2TSB2 (Table 7). Individuals from TSB1TSB1 and TSB3TSB3 carrying *FTA* allele 2 were earlier than individuals from the same group carrying allele 1 (three days for TSB1TSB1 and more than five days for TSB3TSB3, Figure 7). In TSB2TSB2 with individuals carrying allele 1 at *VRN-1A_prom_* headed five days earlier than those carrying allele 2 (Figure 7).

**Table 6:** ANCOVA table with the F statistic and significance for heading date obtained with model (11) and (12). *: *P*_*value*_ < 0.05, **: *P*_*value*_ < 0.01 and ***: *P*_*value*_ < 0.0001, -: not computed in the current model.

| Model | *NBH* | *REP* | *POP* | *GENOGP* | *FTA* | *VRN-1A<sub>ex7</sub>* | *VRN-1D* | *VRN-1A<sub>prom</sub>* | $R^2$ |
| --- | --- | --- | --- | --- | --- | --- | --- | --- | --- |
| (11) | 66.93*** | 0.55 | 7.47*** | - | 1.06 | 2.03 | 0.34 | 26.16*** | 0.37 |
| (12) | 54.47*** | 0.24 | - | 9.83*** | 3.66* | 1.95 | 0.03 | 11.79*** | 0.31 |

**Table 7:**
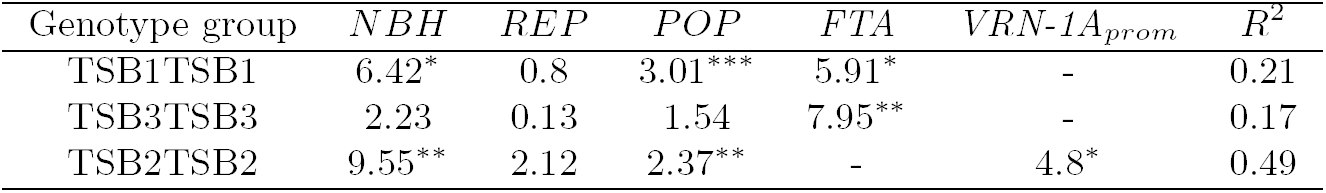
ANCOVA table with the F statistic and significance for heading date obtained with model (11) for TSB1TSB1, TSB2TSB2,TSB2TSB2, the genotype groups with significant effect for at least one of the earliness genes. *: *P*_*value*_ < 0.05, **: *P*_*value*_ < 0.01 and ***: *P*_*value*_ < 0.0001, -: not computed in the current model.

| Genotype group | *NBH* | *REP* | *POP* | *FTA* | *VRN-1A<sub>prom</sub>* | $R^2$ |
| --- | --- | --- | --- | --- | --- | --- |
| TSB1TSB1 | 6.42* | 0.8 | 3.01*** | 5.91* | - | 0.21 |
| TSB3TSB3 | 2.23 | 0.13 | 1.54 | 7.95** | - | 0.17 |
| TSB2TSB2 | 9.55** | 2.12 | 2.37** | - | 4.8* | 0.49 |

**Figure 6:**
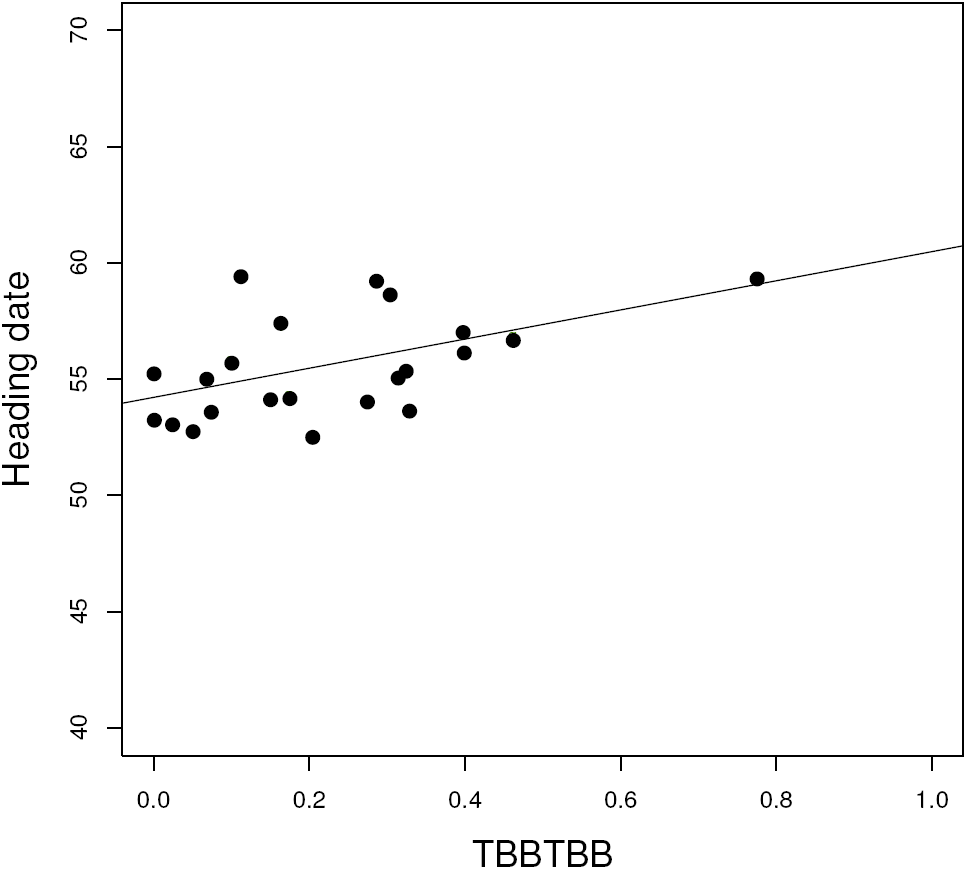
Correlation between LSmean values of heading dates per population and TBBTBB frequency in each population (*R*^2^ = 0.24 and *P*_*value*_ = 0.012).

**Figure 7:**
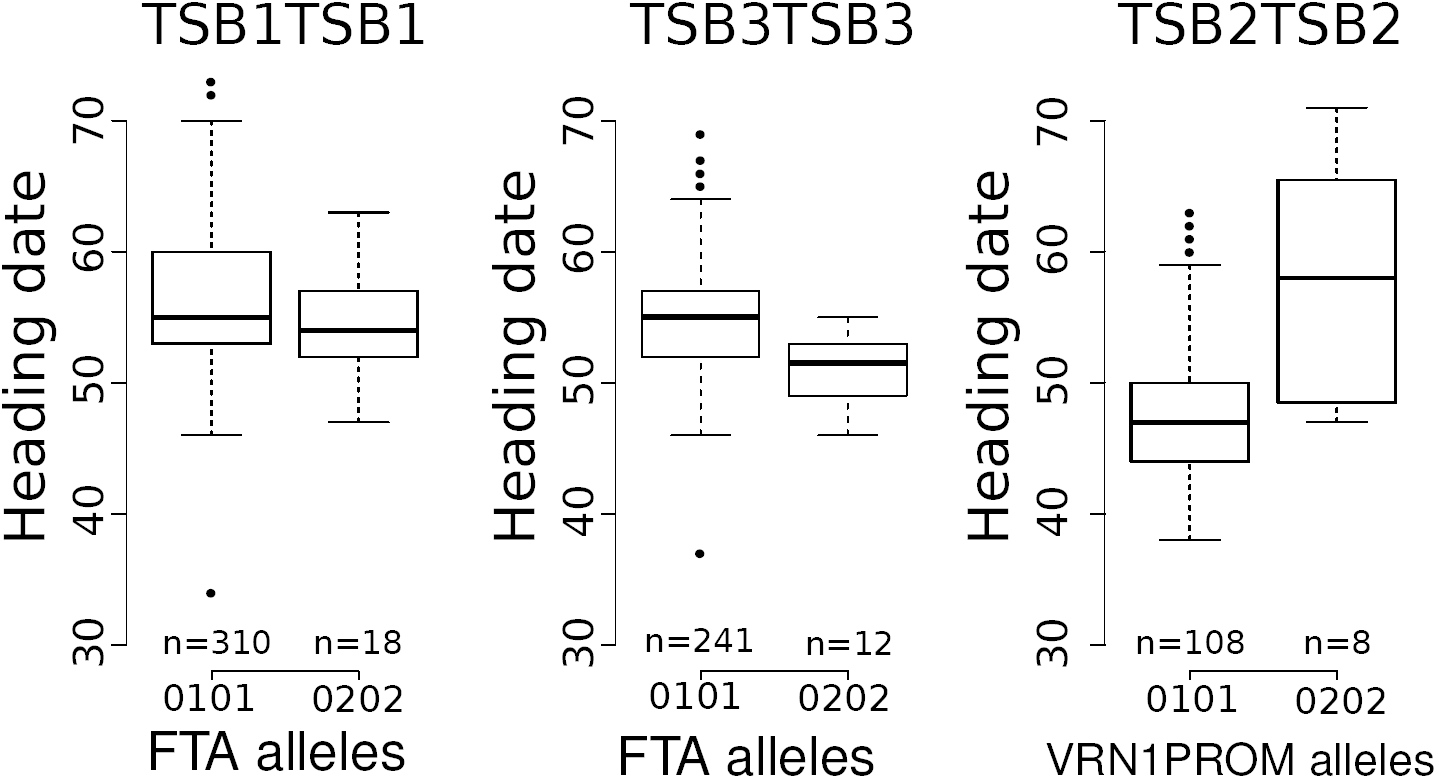
Association between alleles and heading date for three genotype groups (TSB1TSB1, TSB3TSB3 and TSB2TSB2) and two genes (*FTA* and *VRN-1A_prom_*). Class sizes are mentioned under each boxplot.

## 5 Discussion

Mixtures of crop varieties are rather widespread among organic and low-input farmers due to their year in, year out robustness that makes them more stable vis-á-vis variations in biotic and abiotic pressures (Dawson and Goldringer, 2012; Wolfe *et al.*, 2008). However too little attention has been paid to the genetic mechanisms that underlie the micro-evolution of such populations simultaneously submitted to natural and human selection. Studying an on-farm evolutionary experiment allowed us to characterize at the fine genetic level, the short-term spatial and temporal evolution of a mixture of four wheat landraces distributed among farmers in France, The Netherlands and Italy, therefore providing new insights into the behavior of a farmer-led crop metapopulation. The metapopulation showed significant patterns of differentiation at neutral markers, at candidate genes involved in earliness and at earliness, a quantitative adaptive trait. Population differentiation was influenced by combined demographic and selection processes. We hereby focused on the effect of natural selection in particular climatic pressure, in relation to the farmer’s practices. We did not analyse the role of migration process but rather controlled for the effect of migration in differentiation, by detecting the new genotypes potentially introduced by migration. Based on that we defined a subset of the data to detect selection working only with the main genotype groups already present in the initial population (HEF03fld) used as the reference.

### 5.1 Initial structure of the MDT mixture

Among the four components of MDT, TBB, the *Triticum turgidum* landrace and TAN showed a low genetic diversity compared to TSB and TBP, with TBP embedded within TSB. Moreover, the genetic structure of TBP and TSB was similar to that of Rouge de Bordeaux, another on-farm managed population-variety of bread wheat (Thomas *et al.*, 2012). These landraces were composed of a few major haplotypes connected to much less frequent haplotypes in the periphery. These networks of frequent haplotypes connected to numerous rare haplotypes were defined as genotype groups. Relatedness among genotype groups also indicated that they shared part of their genetic background even if they showed specific alleles that allowed us to distinguish among them using control individuals.

In the dynamics of the MDT metapopulation, we considered that a new sub-population appeared when a seed lot was sown on a new farm or on the same farm but with different farming practices. That would correspond to a colonization of an empty patch in the case of natural populations. According to the metapopulation framework, two types of colonization can be considered: the migrant-pool model when an empty patch is colonized by several seed sources and the propagule-pool model when an empty patch is colonized by a unique seed source (Slatkin, 1977). In the case of the MDT mixture, the creation of the mixture by HEF followed the migrant-pool model in a first approximation since each component resulted from independent evolutionary trajectories that have been merged recently. The impact of the founder effect following the migrant-pool model was analysed by comparing the composition of the on-farm reference sample HEF03fld to the virtual MDT. The two populations greatly differed with only 11% of shared haplotypes. This was probably due to differences in the samples provided by CLM to HEF in 1997 and to us in 2008. Indeed, during the multiplication phase of *ex situ* wheat accessions, a limited number of seeds (60) are sown and a strong morphological selection is applied on plants to conserve only one representative spike-type per accession. Moreover, only a few plants that correspond to the “type” are self-pollinated to produce the regenerated sample, leading to strong genetic drift effects. Genetic drift effects have already been observed when within-landrace genetic diversity conserved *ex situ* for different time periods was compared to the diversity conserved on-farm (for barley (Parzies *et al.*, 2000), for bean (Gmez *et al.*, 2005), for pea (Leino *et al.*, 2013)). In addition, when people ask for accessions, the genebank provides them with 40 to 50 seeds per accession. Therefore, a moderate to strong bottleneck effect might occur on the initial level of within-accession genetic diversity, corresponding to the founder effect when the number of individuals involved in the colonization is small (Slatkin, 1977; Wade and McCauley, 1988). HEF received samples of 40-50 seeds from CLM in 1997 for each of the four landraces. In this study, virtual MDT was obtained with seeds provided by CLM after a last multiplication in 2004. Thus, each variety was multiplied from two to three times between 1997 and 2004. Whereas the precise composition of the initial mixture was not available, interviews with farmers and the genebank curator provided some evidences that seed management practices had important demographic consequences at the early stage of the MDT mixture.

### 5.2 Evolution of the MDT mixture under on-farm management

After this first phase of mixture creation, samples of the HEF’s MDT were distributed to different farmers and the MDT populations evolved as independent entities in each particular environment. As a unique source of the mixture was provided to farmers, we considered the diffusion process closer to the propagule-pool model (Slatkin, 1977). We showed that the level of genetic diversity was the same at the metapopulation level (*H*_*E*_ = 0.51) than within the initial population while the haplotype diversity and allele richness were much higher in the metapopulation. Diversity in earliness genes followed the same patterns (Table S1). These results illustrate that submitting several sub-populations to contrasted environmental conditions allowed a good maintenance of the initial allelic diversity at the metapopulation level, showing that crop metapopulations maintain agrobiodiversity in agroecosystems, as expected from the theory (Olivieri *et al.*, 1995). Such a phenomenon was empirically observed for wild inbred populations of *Leavenrworthia* (Liu *et al.*, 1998), bread wheat populations maintained in a dynamic management design (Goldringer *et al.*, 2006) and it was demonstrated theoretically in the context of the metapopulation theory (Ingvarsson, 2002). It has been shown that in the absence of turnover (extinction/recolonization), the increase in genetic differentiation associated with the low pollen migration rates in highly selfing species are the main drivers of the stabilization of the genetic diversity at the metapopulation level.

In the context of propagule-model, differentiation is expected to be particularly high for neutral markers (Pannell and Charlesworth, 1999) and could partly explain the observed pattern of differentiation among populations (0.11) for multilocus *F*_*ST*_, and (0.15) for genotype group *F*_*ST*_. The impact of on-farm management practices will then greatly depend on whether farmers grow their populations in small plots (collection) or in fields, i.e. on population size. Genetic drift has already been reported for wheat and for maize when farmers grew populations in small plots (Zhang *et al.*, 2006; van Heerwaarden *et al.*, 2010). For one farmer (BER) who continuously grew MDT in very small plots (around 80 plants in BER08col), founder effect and genetic drift were combined, leading to one of the lowest genetic diversity (*H*_*E*_ = 0.35) and effective population size (*N*_*E*_ = 10). These hypotheses could be rigorously tested using the theoretical framework developed by van Heerwaarden *et al.* (2010); Artoisenet and Minsart (2014). However, that would require to adapt the model to integrate founder effect for better accounting for farmer practices and social organization.

Differentiation within the metapopulation was assessed at three levels: (i) neutral markers, (ii) polymorphisms in genes associated to earliness and (iii) earliness at the phenotypic level. We focused on earliness for its important role in adaptation to the environmental condition, in particular to the interaction between sowing date (farming practice) and climate. Indeed, populations need to synchronize their reproductive cycle to the climate conditions (Rhoné *et al.*, 2008) which are different from one farm to another and depending on farming practices. Differentiation within the MDT metapopulation was higher at candidate gene (*F*_*STQ*_ = 0.24) and for earliness (*Q*_*ST*_ = 0.26) than at multilocus *F*_*ST*_ = 0.11. This particular pattern of differentiation: *F*_*ST*_ < *F*_*STQ*_ ≤ *Q*_*ST*_, was interpreted as the result of divergent selection (Merila and Crnokrak, 2001; McKay and Latta, 2002), given that there was no or a limited gene flow among the populations (Le Corre and Kremer, 2012). This interpretation relying on theoretical study is consistent with the history of MDT. Different sub-populations of MDT were disseminated in different socio-climatic environments through seed exchanges, leading to quite strong divergent selective pressures on the mixture. In addition, farmer practices aimed at limiting gene flow among crop populations. Local exceptions were observed but did not seem to affect the global pattern of differentiation. A similar pattern was observed for maize populations grown in Mexico (Pressoir and Berthaud, 2003b), although the underlying social and evolutionary processes are different mainly due to the different mating system. Moreover in the MDT metapopulation, the stronger differentiation in the candidate genes showed that 10 of the 15 populations have faced different climatic and farming conditions inducing divergent selection in gene regions involved in adaptive traits such as earliness.

The evolution of the crop metapopulation is mediated by the interaction between farmer-led and natural selection. In the case of MDT management by a farmers’ network, direct farmer-led selection was rare. However, particular practices that were specific to some farmers could have reinforced the effect of natural selection, such as late sowing or harvesting before the seeds of all genotypes were mature. It would be interesting to study whether other fitness-related traits were also differentiated among populations.

### 5.3 Genetic mechanisms involved in earliness differentiation

In the context of a self-pollinated species, selection could significantly affect gene diversity and also neutral diversity in particular around the genes submitted to selection, through hitch-hiking effects and/or background selection (Ingvarsson, 2002). The strong variation in the composition of the mixture was positively correlated with altitude (Figure S2 in supporting information). The specific environmental conditions occurring high up in the mountains might have affected the viability of some particular landraces. *Touselles* landraces are not expected to be adapted to mountain conditions since they historically were grown in southeastern France. Thus, we noticed that TBB, TAN and TBP, the three landraces that were the least diversified (Table 2) consistently decreased in frequency at high altitudes while TSB increased. TSB was the only landrace component that showed a level of diversity (*H*_*E*_ = 0.26) similar to the rare measures available for other wheat landraces from Oman, Turkey and Mexico that ranged between 0.15 and 0.55 (Dreisigacker *et al.*, 2005; Zhang *et al.*, 2006), while TAN, TBB and TBP had a lower level of diversity. Here the local adaptation of the mixture could be the result of two different processes shaping the standing variability available within the mixture: (i) selection among components of the mixture; ii) selection within the components. These two processes are discussed hereby.

A strong differentiation among genetic group was found for earliness. Thus, we assumed that change in mixture composition could affect the average earliness value (Figure 5 and Table 4). The positive correlation between the frequency of TBBTBB and the heading date of populations suggested that at least part of the phenotypic divergence was due to variation in the proportion of TBBTBB.

In addition to the evolution of the mixture composition, two landraces (TBB and TSB) showed significant among-population divergence for earliness, indicating a within-landrace genetic evolution.

In order to understand more finely the genetic basis of earliness differentiation we studied genes known to be associated with earliness. All landraces had a very low level of genetic diversity for the earliness gene polymorphisms (*H*_*e*_ between 0 and 0.03, Table S1). Allele fixation at earliness genes in these landraces induced statistical confusion between the effects of the genotype group and of the gene. This could explain the lack of association between earliness and polymorphism at two of the four earliness genes in these populations whereas a strong association was detected in a core collection of bread wheat (Rousset *et al.*, 2011). Association between *FTA* and *VRN-1A_prom_* polymorphisms and earliness was detected within the TSB groups present in the metapopulation of MDT (Figure 7, Table 5). Therefore, the genetic evolution within landraces in the different mixtures seems to play a significant role in the differentiation of the populations for heading date. This is consistent with the fact that TSB was the most frequent in many populations and was maintained at a very high level in the metapopulation indicating a higher adaptability. Moreover, such a body of evidence indicates that TSB still kept the ability to adapt to contrasted environments. Specific additional experiments would be needed to confirm this hypothesis.

In addition, we found that different genetic trajectories led to the same heading date at the population level as illustrated in Figure 8. For instance, two populations were late heading with a high frequency of TBBTBB (OLR08col) but also with a very low frequency of TBBTBB (RAB08col). Paradoxically, the same frequency of TBBTBB and TSB1TSB1 in another population was associated to an early heading date (BER08col, light color). These findings provide an example of the strong genetic “plasticity” of mixtures and their ability to adapt to different environments. In spite of the challenges posed by the on-farm evolutionary experiment where populations are submitted to real farming conditions, we were able to highlight how the standing genetic variation is important in adaptive processes.

**Figure 8:**
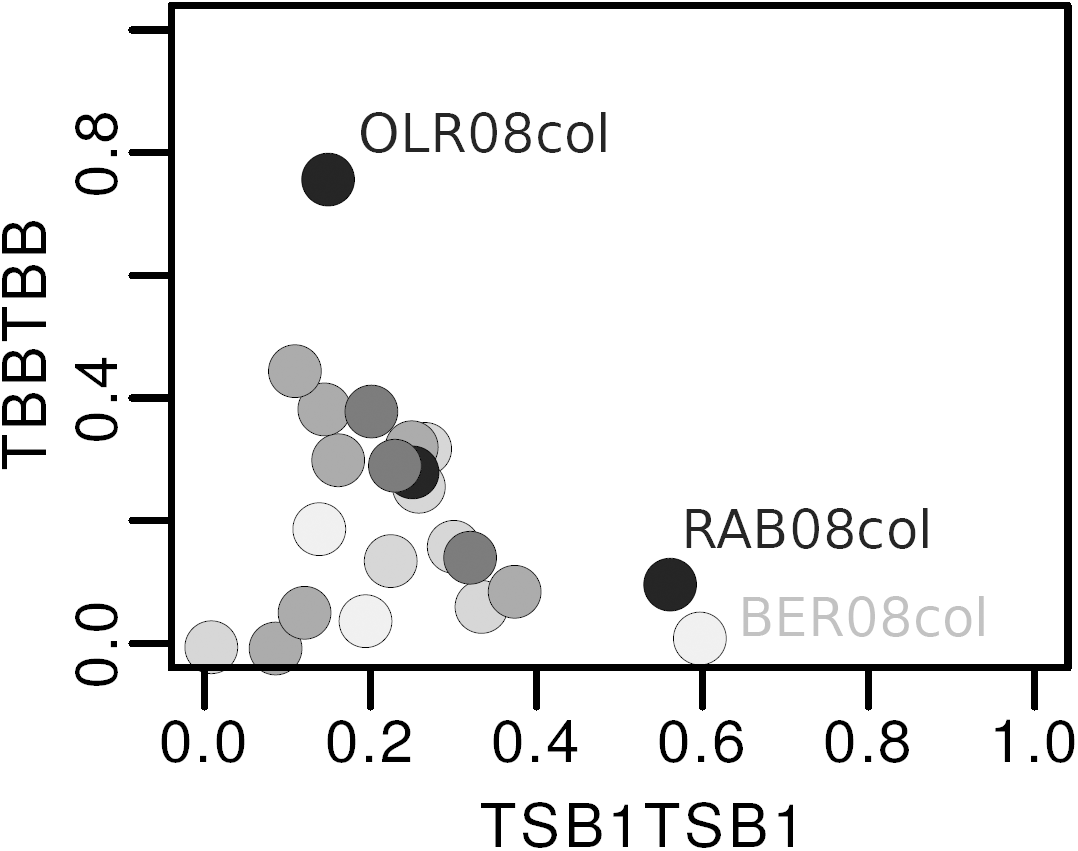
Effect of TSB1TSB1 and TBBTBB frequencies on the mean heading date per population. Heading date values are represented by the different colors. Dark colors represent late heading dates, light colors represent early heading dates

## 6 Conclusion

This paper aimed to depict the main genetic characteristics of a recently established crop mixture, and its evolution within a farmer-led seed exchange network, considering this particular design of On-Farm Dynamic Management (OFDM) as an on-farm evolutionary experiment. Submitted to contrasted environments and practices, within metapopulation genetic diversity was maintained over time, while multi-level differentiation among populations was detected. Particular agricultural practices were identified as playing an important role in genetic drift or in selection, leading to a specific differentiation pattern. Moreover, natural selection associated to the contrasted farming practices shaped the genetic and phenotypic divergence of the populations through the diversity of the environments where the mixture evolved.

Our findings highlighted the remarkable ability of the mixture to respond to selection in drastic conditions. While we initially expected that population differentiation would be mostly mediated through variation in the proportions of the mixture components, we found that within component genetic evolution also substantially contributed. In particular, the TSB landrace, the most diversified landrace of the four was identified as the keystone in the adaptation process of the mixture. This landrace was present in all populations and it responded with different strategies for earliness depending on global environmental and agricultural conditions. These findings emphasize how critical it is to maintain within-variety genetic diversity. The distribution of crop genetic diversity met in OFDM is a product of the self-organization of the farmers and therefore this study showed that such social organization contributes to the adaptation of crop biodiversity to climate change. In addition, this short term evolutionary experiment sets the stage for promising properties of mixtures and confirms the potential of genetic diversity to maintain adaptability and stability in changing environments. This investigation needs to be continued through the medium term to confirm these results.

## Acknowledgments

The authors gratefully acknowledge Élise Demeulenaere for her advises during the survey preparation. The authors thank Audrey Didier from the Clermont-Ferrand genebank for providing *ex situ* accessions, the farmers of the Réseau Semences Paysannes for stimulating discussions. This work was supported by a grant from the Institut National de la Recherche Agronomic, France provided to M. T. A grant from the Bureau des Ressources Génétiques (now Fondation sur la Recherche sur la Biodiversité) (2005-2007 project) provided money for field and molecular experiments.

